# Genome-wide rules of nucleosome phasing

**DOI:** 10.1101/093666

**Authors:** Sandro Baldi, Dhawal S. Jain, Lisa Harpprecht, Angelika Zabel, Marion Scheibe, Falk Butter, Tobias Straub, Peter B. Becker

## Abstract

Regular successions of positioned nucleosomes – phased nucleosome arrays (PNAs) – are predominantly known from transcriptional start sites (TSS). It is unclear whether PNAs occur elsewhere in the genome. To generate a comprehensive inventory of PNAs for *Drosophila*, we applied spectral analysis to nucleosome maps and identified thousands of PNAs throughout the genome. About half of them are not near TSS and strongly enriched for a novel sequence motif. Through genome-wide reconstitution of physiological chromatin in *Drosophila* embryo extracts we uncovered the molecular basis of PNA formation. We identified Phaser, an unstudied zinc finger protein that positions nucleosomes flanking the new motif. It also revealed how the global activity of the chromatin remodeler CHRAC/ACF, together with local barrier elements, generates islands of regular phasing throughout the genome. Our work demonstrates the potential of chromatin assembly by embryo extracts as a powerful tool to reconstitute chromatin features on a global scale *in vitro.*

## Introduction

DNA in the eukaryotic nucleus is stored as chromatin, consisting of repeating nucleosome units, in which 147 bp segments of DNA are wrapped around histone octamers (Luger et al., 1997). Nucleosomes are not only important for compacting and storing DNA in the nucleus, but they are critically involved in regulating the accessibility of the DNA to polymerases and protein regulators of RNA transcription, DNA replication and damage repair (Groth et al., 2007; Radman-Livaja and Rando, 2010; Struhl and Segal, 2013). The exact position of nucleosomes relative to the underlying DNA sequence – referred to as nucleosome positioning – has important consequences for the recognition of sequence motifs on DNA (Radman-Livaja and Rando, 2010; Struhl and Segal, 2013; Zaret and Carroll, 2011). If several adjacent nucleosomes are well-positioned, they are said to form a phased nucleosome array. Phased regular arrays of nucleosomes, i.e. stretches of well-positioned nucleosomes with an even spacing between them, are best known downstream of active transcriptional start sites (TSS) (Mavrich et al., 2008; Rube and Song, 2014; Teif et al., 2012; Valouev et al., 2011; Yuan et al., 2005). There, it is thought that the +1 nucleosome immediately downstream of the TSS is strongly positioned by the adjacent nucleosome-free region, the binding of general transcription factors and the action of ATP-dependent chromatin remodelers (Hartley and Madhani, 2009; Krietenstein et al., 2016; Lieleg et al., 2015; Struhl and Segal, 2013; Zhang et al., 2011). The nucleosomes further downstream are regularly phased relative to the +1 nucleosomes by chromatin remodelers and possibly transcriptional elongation (Gkikopoulos et al., 2011a; Krietenstein et al., 2016; Lieleg et al., 2015; Yen et al., 2012; Zhang et al., 2011). A second known case of regular phasing of nucleosome arrays occurs adjacent to binding sites of the insulator protein CTCF (Fu et al., 2008; Wiechens et al., 2016).

Are there any other nucleosome phasing elements in the genome? And what does it take to generate them? Current descriptions of phased nucleosome arrays are commonly based on cumulative average nucleosome profiles and therefore bound to reference DNA sequence features such as TSS or CTCF binding sites. This restricts the analysis to well annotated candidate regions in the genome. Furthermore, if nucleosome positioning relative to the alignment point is not the same at most averaged sites, regular phasing will not be apparent in the averaged plot, even if each single site is regularly phased itself.

To overcome these shortcomings and systematically survey a genome for regularly phased nucleosome arrays we developed two new tools and applied them to the genome of *Drosophila melanogaster:* (1) we mapped regular, phased nucleosome arrays in an unbiased manner by spectral density estimation and (2) we reconstituted complex chromatin at the genomic scale in a cell-free system, allowing a mechanistic dissection of nucleosome phasing. Interestingly, our comprehensive map of phased regular nucleosome arrays revealed that only half of them coincide with TSS. In addition, we found several new classes of potential barrier elements, i.e. DNA sequences that serve as barriers for extended nucleosome phasing. Several of these barrier elements corresponded to various low-complexity repeat sequences. Only one class corresponded to the binding site for a known chromatin protein, the insulator protein ‘suppressor of hairy wing’ [su(Hw)]. The most abundant barrier element was characterized by a novel sequence motif, which is bound by ‘Phaser’, a previously uncharacterized zinc finger protein. Finally, we show that extensive nucleosome phasing at Phaser and su(Hw) sites is achieved in cooperation between local barrier elements and the global activity of the CHRAC/ACF chromatin remodeler.

## Results

### A comprehensive inventory of phased regular nucleosomes throughout the genome

To identify regions of regularly phased nucleosomes throughout the genome, we generated genome-wide nucleosome dyad maps in 2-8 h old *D. melanogaster* embryos in five replicates (three in the *w1118* and two in the *yw* background) (**Fig. 1A**). The regularity of nucleosome arrays can be best appreciated if they are phased by alignment to a DNA sequence or protein barrier that operates in all cells, as, for example, the nucleosome-free region at TSS of housekeeping genes (Lieleg et al., 2015; Radman-Livaja and Rando, 2010; Struhl and Segal, 2013). However, such an alignment requires prior knowledge of candidate phasing sequences. To survey the genome for phased nucleosomes in a more unbiased manner, we applied spectral density estimation (SDE) to sliding 1 kb windows along the nucleosome dyad density profiles (**Fig. 1A**). This method treats nucleosome occupancy as a composite periodic signal and decomposes it into a linear combination of simple periodic functions using Fourier transformation. The weights (or spectral densities) and periods of the decomposed functions determine the overall contribution of a given period to the composite pattern. The heat map indicates that sites of regularly phased nucleosome arrays are revealed by the weight of a periodicity corresponding to a nucleosome density of roughly 5-6 nucleosomes per 1000 bp. Therefore, we looked at the spectral density corresponding to a period of 192 bp (5.2 nucleosomes per 1kb). The strength of the SDE at each genomic position then indicates how much the nucleosomes in the surrounding 1 kb resemble a completely regular array with 192 bp periodicity. By assigning an empirically determined threshold, we identify 10417 regions of high nucleosome array regularity throughout the genome (**Fig. 1A, blue shaded**). We call these sites phased regular nucleosome arrays (PNA). Their median length is 700 bp, accounting for 6.2% of the euchromatic part of the genome. PNAs are enriched 4.1 fold at promoters (**Fig. 1B**), which is expected since nucleosomes adjacent to many active promoters are regularly phased (Mavrich et al., 2008; Rube and Song, 2014; Teif et al., 2012; Valouev et al., 2011; Yuan et al., 2005). This was also apparent when we determined the distribution of PNAs relative to the 9-chromatin-states model (Kharchenko et al., 2011), which showed a 4.3 fold enrichment in ‘state 1’ chromatin (active promoters) (**Fig. 1C**).

**Fig. 1.**
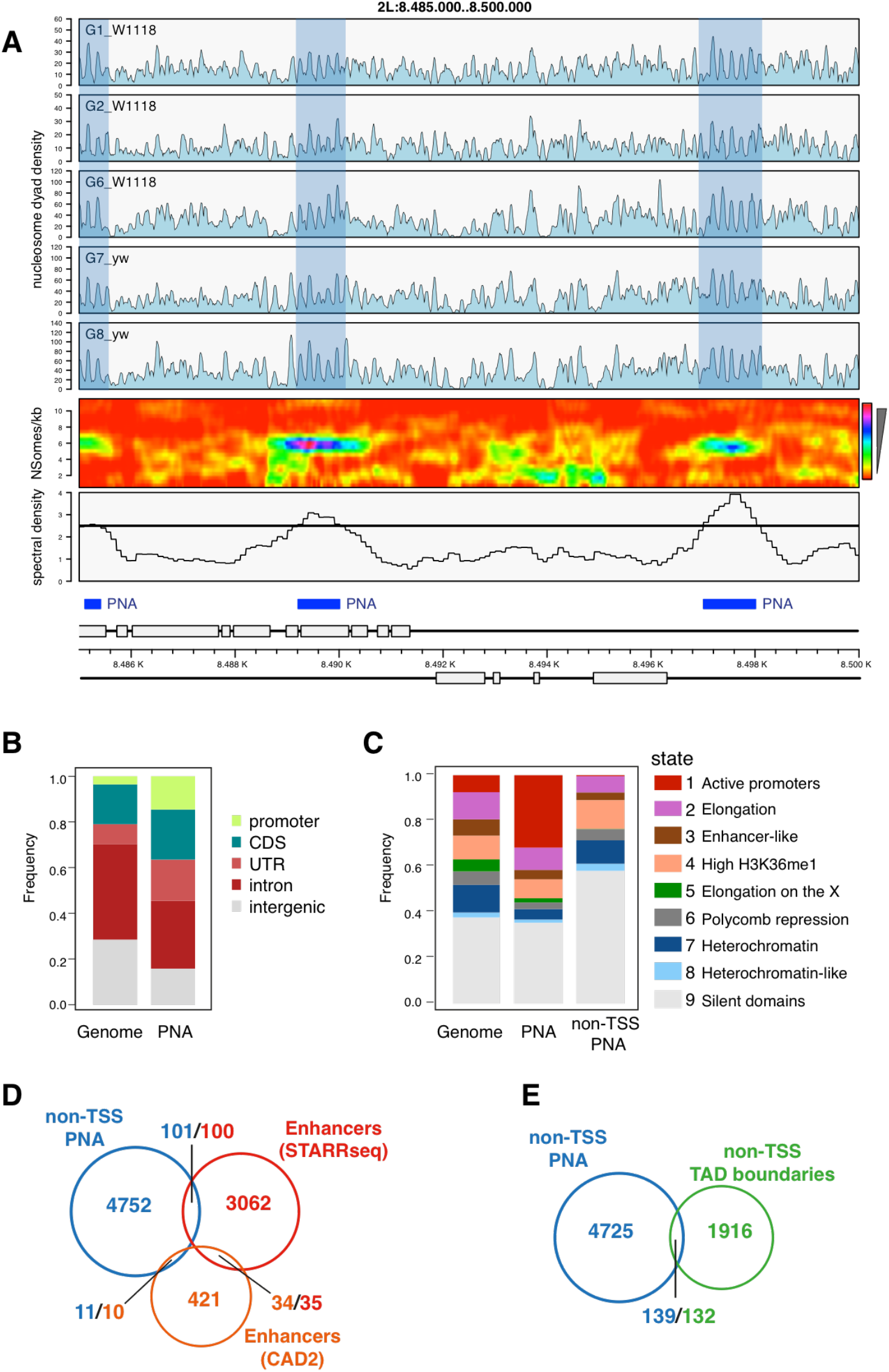
A comprehensive inventory of phased regular nucleosomes throughout the genome. (A) Nucleosome dyad densities for a representative genomic region for 5 *WT* samples of two different genotypes (upper panels). Heat map: spectral densities obtained from sample “G8_yw” are displayed for the range of 1-11 nucleosomes per kb (periods of 100-500 bp). Lower panel: Spectral density corresponding to a frequency of 5.2 nucleosomes per 1 kb (period of 192 bp) after standardization and averaging across all the samples. An empirical threshold set at 2.5 defines phased nucleosome arrays (PNA, blue boxes and shades). (B) Genomic feature annotation of PNA regions (n=10417). (c) Chromatin state distribution of PNA (n=10417) and PNA that are distal to promoters (non-TSS PNA, n=4864). (D) Overlap of non-TSS PNA with annotated enhancer regions. Two different numbers in the overlaps indicate that single features from one category may overlap with multiple features from another one. (E) Overlap of non-TSS PNA with TAD boundaries that are themselves not at TSS. The two different numbers in the overlap indicate that single TAD boundaries may overlap with multiple non-TSS PNAs.

Interestingly, almost half of all PNAs (4864/10417) are actually not close to any promoter. We call them ‘non-TSS PNA’ and decided to study them in more detail. This group was strongly depleted of ‘state 1’ chromatin (active promoters) but enriched in ‘state 9’ (silent domains) (**Fig. 1C**). Interestingly, there was only a poor overlap with known enhancer sites (Arnold et al., 2013; Bonn et al., 2012) (**Fig. 1D**) and TAD boundaries (Ramirez et al., 2018) (**Fig. 1E**). Using the ChIP-Atlas tool (Oki et al., 2018) to identify chromatin immunoprecipitation (ChIP) profiles that would resemble the distribution of non-TSS PNA, we did not find any enriched histone modifications at these sites. However, the query revealed that multiple insulator proteins, and particularly the *gypsy* insulator subunit su(Hw), are enriched at non-TSS PNA (Supp. Table 1).

### A new DNA motif is strongly enriched at non-TSS phased regular nucleosome arrays

To explore whether non-TSS PNA shared any common sequence feature, we applied MEME (Bailey, 2011) for *de novo* motif identification to a subset of non-TSS PNA that were at least 1 kb long (corresponding to minimally 5-6 phased nucleosomes). This revealed a list of 10 enriched motifs (**Fig. 2A, PPM in Supp. Table 2**). The most highly enriched motif is a complex signature characterized by a central ATACG motif, which, interestingly, shows no similarity to known motifs in the JASPAR database (Mathelier et al., 2014). Almost a third of non-TSS PNA contained this motif, suggesting it might be a major determinant of nucleosome phasing outside of promoters. ‘Motif 5’ is very similar to a published consensus site for the insulator protein su(Hw) (Adryan et al., 2007; Baxley et al., 2017), which fits to our observation that su(Hw) binding profiles resemble the distribution of non-TSS PNA. The remaining motifs were predominantly signatures of relatively low complexity.

**Fig. 2.**
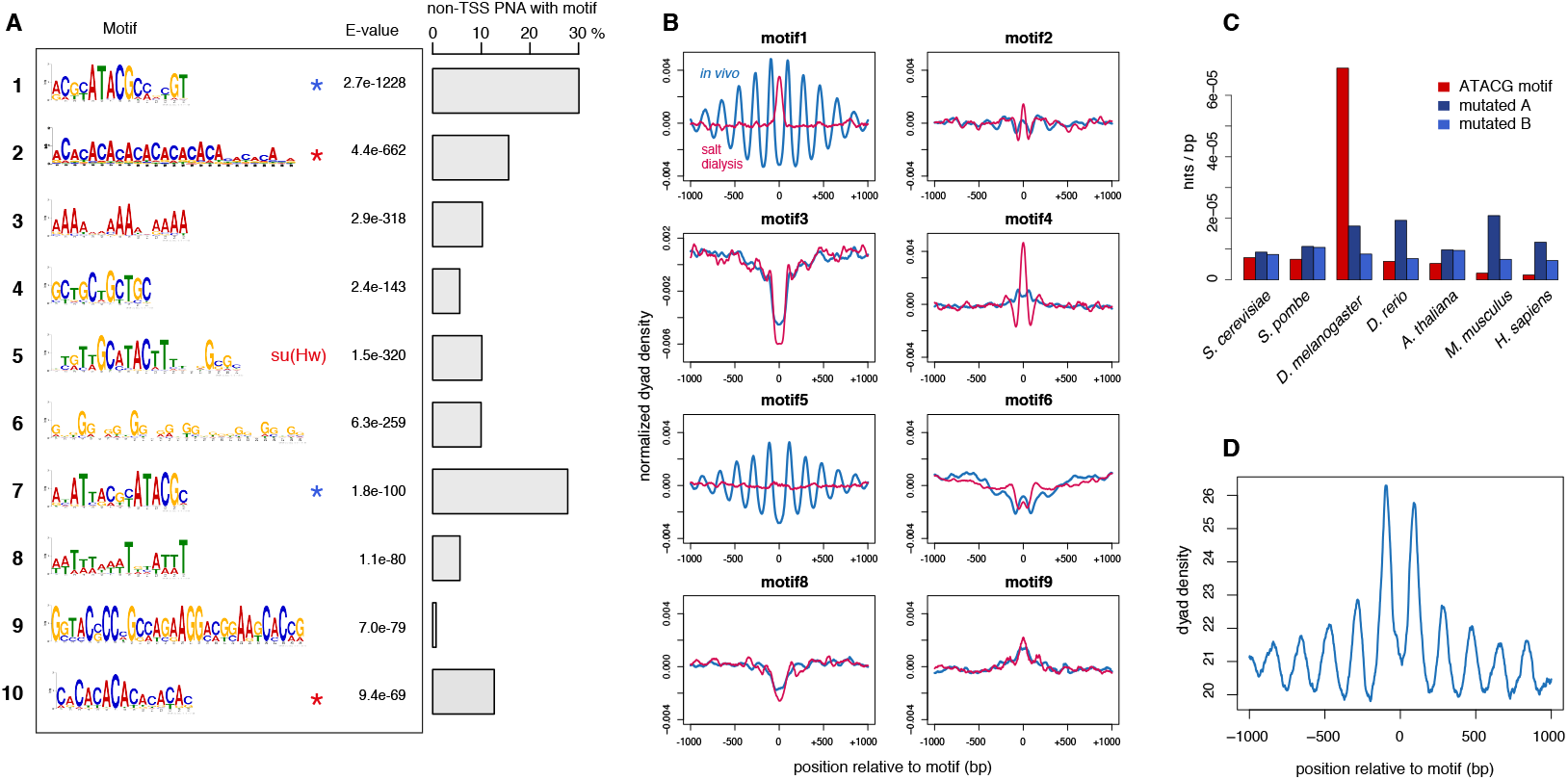
A new DNA motif is strongly enriched at non-TSS phased regular nucleosome arrays. (A) Motifs enriched at non-TSS PNA. Equivalent motifs (#1#/7, #2/#10) are indicated by colored asterisks. Motif 5 closely resembles the consensus binding motif for the insulator protein su(Hw). On the right: percentage of non-TSS containing a particular motif. (B) Nucleosome dyad densities aligned at highly significant motif sites throughout the genome in 2-8h old *Drosophila* embryos (blue) and in chromatin assembled by salt gradient dialysis of purified histones on genomic *Drosophila* DNA (red). (C) Occurrence of the ATACG motif (#1) in different species compared to two related control motifs. (D) Nucleosome dyad densities aligned at 4295 ATACG consensus sites in the mouse genome.

For each motif, we determined highly significant consensus sites throughout the genome and used them to align nucleosome dyad profiles from 2-8 h old embryos (**Fig. 2B**, blue lines). At the ATACG motif and the putative su(Hw) sites, a pattern of 8-10 regularly phased and well-positioned nucleosomes emerged, with the motifs directly placed in the central nucleosome linker. Interestingly, while the central nucleosome linkers that harbor the su(Hw) sites appear expanded as if the bound protein (presumably the *gypsy* insulator complex) left a ‘footprint’ in the nucleosome array, such an expansion was not observed for linkers containing the ATACG motif. We also observe high chromatin accessibility in ATAC-seq profiles around su(Hw) binding sites but not at ATACG motifs (**Supp. Fig. 1**). The other motifs did not show any obvious regular phasing on the averaged profiles. Conceivably, those low-complexity sequence elements are characterized by length heterogeneity at different genomic loci. This would explain why they are picked up by SDE at individual sites, but do not lead to phased arrays in the cumulative plots.

Nucleosome positioning along DNA depends on the underlying DNA sequence and its propensity to form nucleosomes, but also on DNA binding proteins and chromatin remodelers (Lieleg et al., 2015; Radman-Livaja and Rando, 2010; Struhl and Segal, 2013). To determine the intrinsic nucleosome positioning potential of the identified motifs in their genomic context and in the absence of any *trans*-acting factors, we assembled nucleosomes from purified histones by salt gradient dialysis (SGD) on total genomic DNA and mapped the nucleosomes by MNase-seq (**Fig. 2B**, red lines). This revealed that nucleosome positioning at ATACG and su(Hw) sites is not an intrinsic property of DNA, as SGD-reconstituted nucleosomes are not excluded from ATACG or su(Hw) sites (**Fig. 2B**). Remarkably, SGD-nucleosomes were even preferentially assembled on the ATACG site, suggesting active nucleosome exclusion at these sites *in vivo.* Nucleosome positions at motifs 2, 3, 6, 8 and 9 was similar *in vivo* and in SGD chromatin, suggesting that positioning at these sites may be predominantly determined by DNA-intrinsic features. This is evident for motifs 3 and 8, which are A/T rich, a feature that disfavors the formation of nucleosomes (Nelson et al., 1987). Accordingly, these sites are devoid of nucleosomes and presumably flanked by well-positioned nucleosomes at the level of single sites.

### The association of the ATACG motif with regularly phased nucleosomes is conserved in mammals

With the ATACG sequence motif we identified a possible new and strong determinant of phased regular nucleosome arrays throughout the genome. Interestingly, the occurrence of this motif in the *D. melanogaster* genome is much more frequent than the occurrence of two related motifs with nucleotide substitutions in the consensus sequence (**Fig. 2C**), indicating there is a positive selection for it. On the other hand, the same motif appears to be depleted from the mouse and human genomes relative to control motifs, whereas other species do not show any strong bias. The apparent counter-selection of the motif in mammals suggested that it might serve a conserved function. To investigate nucleosome pattering around the motif in mammals, we identified 4295 ATACG consensus sites in the mouse genome. Aligning nucleosome dyad densities from high-quality mononucleosome profiles (Voong et al., 2016) to these sites yielded a very similar pattern as in *Drosophila*, with two well-positioned nucleosomes flanking the motif and regularly phased nucleosomes on either side (**Fig. 2D**). On the other hand, aligning the same data set to su(Hw) consensus sites in the mouse genome did not reveal any strong nucleosome positioning or regularity (**Supp. Fig. 2A**).

### The ATACG motif is required for the formation of phased nucleosome arrays *in vitro*

We had established a strong correlation between the presence of the ATACG motif and the formation of PNAs throughout the genome. Next, we wanted to assess whether the motif is also causal to the formation of such arrays. To this end, we developed a new strategy that renders nucleosome positioning and protein-chromatin interactions in the *Drosophila* genome accessible to biochemical analysis. Extracts of 0-90 min old preblastoderm *Drosophila* embryos (DREX) are rich in histones, chaperones, remodelers and many other chromatin factors. These extracts form the basis of an efficient cell-free assembly system for complex chromatin on DNA (Becker and Wu, 1992; Volker-Albert et al., 2016). So far, the system had only been used to assemble chromatin on bacterial plasmid DNA. To reconstitute the entire fly genome into chromatin, we purified long (~50 kb) fragments of genomic DNA from *Drosophila* BG3-c2 cells and incubated them with DREX and an ATP-regenerating system. The *in vitro* assembled chromatin was then digested with MNase, and the mononucleosomal DNA fragments were sequenced and mapped to the fly genome (**Fig. 3A**). When the nucleosome dyad densities were aligned to 1737 ATACG sites in the fly genome, the *in vitro* pattern was very similar to the one obtained from embryos *in vivo* (**Fig. 3B**). The cell-free system also reconstituted PNAs adjacent to ‘motif 5’, the putative su(Hw) sites. On the other hand, nucleosome phasing at TSS was only rudimentarily reconstituted *in vitro*, not surprisingly, since extracts from preblastoderm embryos are not transcriptionally active (Becker and Wu, 1992). We also determined PNAs by spectral density estimation on genome-wide nucleosome maps of chromatin assembled *in vitro.*

**Fig. 3.**
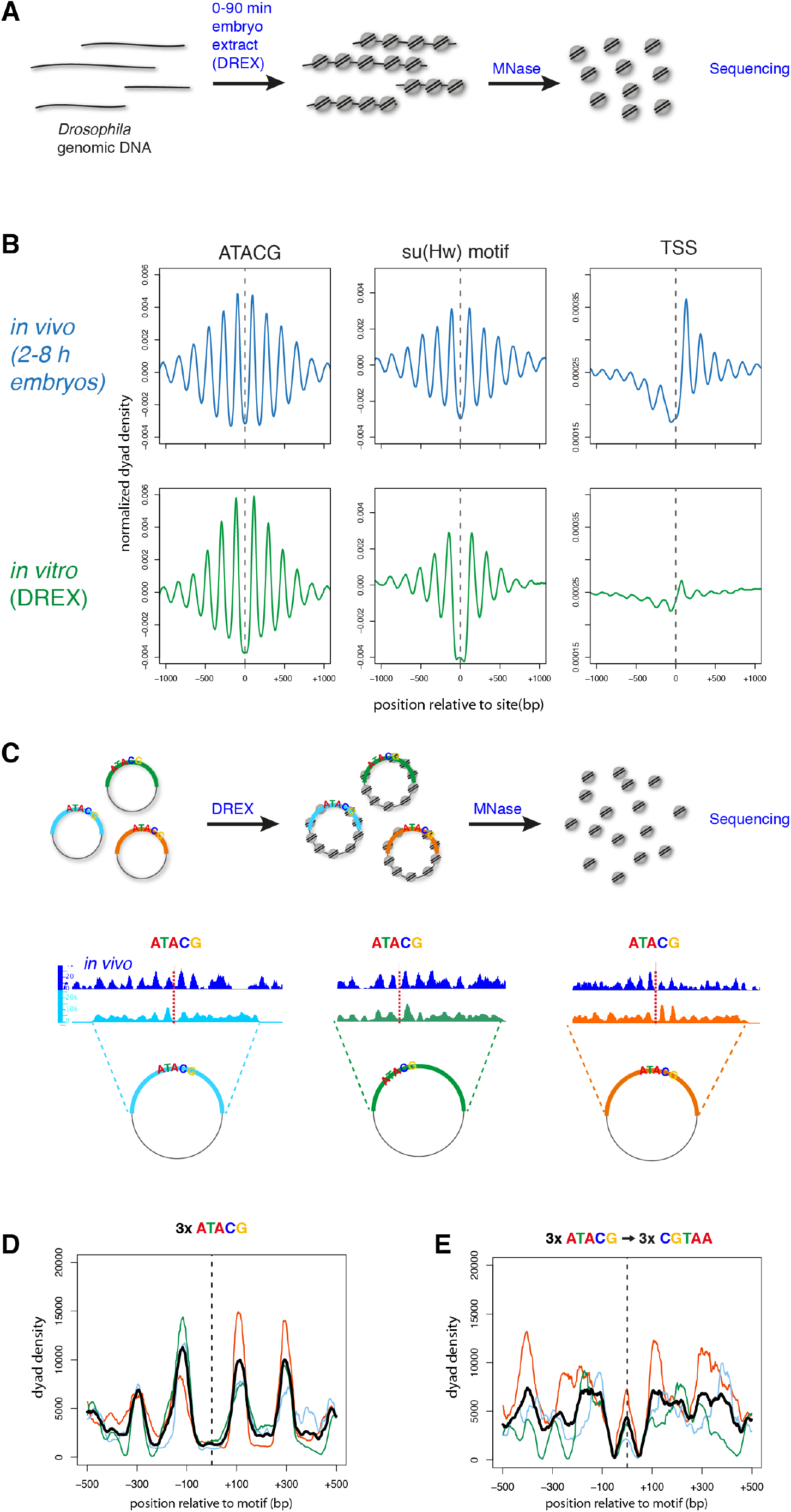
The ATACG motif is required for the formation of phased nucleosome arrays *in vitro*. (A) Chromatin is assembled *in vitro* on a genome-wide scale by incubating *Drosophila* genomic DNA in *Drosophila* preblastoderm embryo extract (DREX). Nucleosome positions are determined by MNase-digestion and sequencing of mononucleosomal fragments. (B) Averaged nucleosomal dyad densities aligned at ATACG-sites, su(Hw) motifs and TSS *in vivo* (2-8 h embryos, blue) and *in vitro* (DREX, green). (C) Three PNAs containing ATACG motifs were cloned into plasmids and used for chromatin assembly by DREX. Sequencing of mononucleosomal fragments reveals regularly phased nucleosomes around the ATACG motifs. (D) Nucleosomal dyad densities from (C) aligned at the ATACG motifs. Averaged density in black. (E) Cloned PNAs were mutated in the ATACG motifs and used for *in vitro* chromatin assembly and nucleosome mapping. Shown are nucleosomal dyad densities aligned at the mutated ATACG motifs and the averaged density in black.

*De novo* motif identification on all *in vitro* PNAs yielded the ATACG motif as the top hit (**Supp. Fig. 2B**), establishing it as the major marker of local nucleosome phasing in the absence of transcription. Thus, we concluded that the faithful *in vitro* reconstitution of nucleosome phasing at ATACG sites on a global scale renders the phenomenon amenable to further mechanistic dissection on single sites.

To determine whether the ATACG motif was indeed important for nucleosome phasing, we cloned three different genomic sites along with 2 kb of flanking sequence which are decorated with well-defined, regularly phased nucleosomes *in vivo.* These three plasmids were assembled into chromatin *in vitro* and the nucleosomes mapped (**Fig. 3C**). Aligning the nucleosome tracks from the three cloned regions at the motif sites revealed a similar pattern as *in vivo* with two well-positioned nucleosomes flanking the motif (**Fig. 3D**). Mutating the central five nucleotides of the ATACG motif on the plasmids from ATACG to CGTAA was sufficient to largely abolish regular phasing of nucleosomes around the sites and even led to nucleosome formation on the mutated site (**Fig. 3E**). Together, these findings reveal a causal role for the ATACG motif in the formation of PNAs.

### Identification of the ATACG-binding protein Phaser

We found the ATACG motif to be associated with many non-TSS PNAs and required for the formation of phased arrays. However, we have also shown that nucleosome positioning around ATACG sites cannot be explained by intrinsic DNA properties alone. Thus, we wished to purify any protein that would bind to the motif and may serve as a barrier factor to set up PNAs throughout the genome. Our strategy was informed by two observations: (1) The factor should be present in preblastoderm embryo extract, since *in vitro* chromatin assembly faithfully recapitulated nucleosome phasing around ATACG sites. (2) Conceivably, an ATACG-binding protein may physically interact with the flanking nucleosomes, which may contribute to their positioning, but also stabilize ATACG binding. Therefore, we set out to purify the factor bound to a linker of a dinucleosome reflecting its native environment. To do this, we cloned a genomic dinucleosome-sized DNA fragment featuring an ATACG motif in the central linker. Oligomerizing the fragment and coupling to paramagnetic beads created an affinity reagent. We reasoned that incubation of the bead-bound DNA under chromatin assembly conditions should reconstitute nucleosome positioning and ATACG binding in every second nucleosome linker (Fig. 4A).

**Fig. 4.**
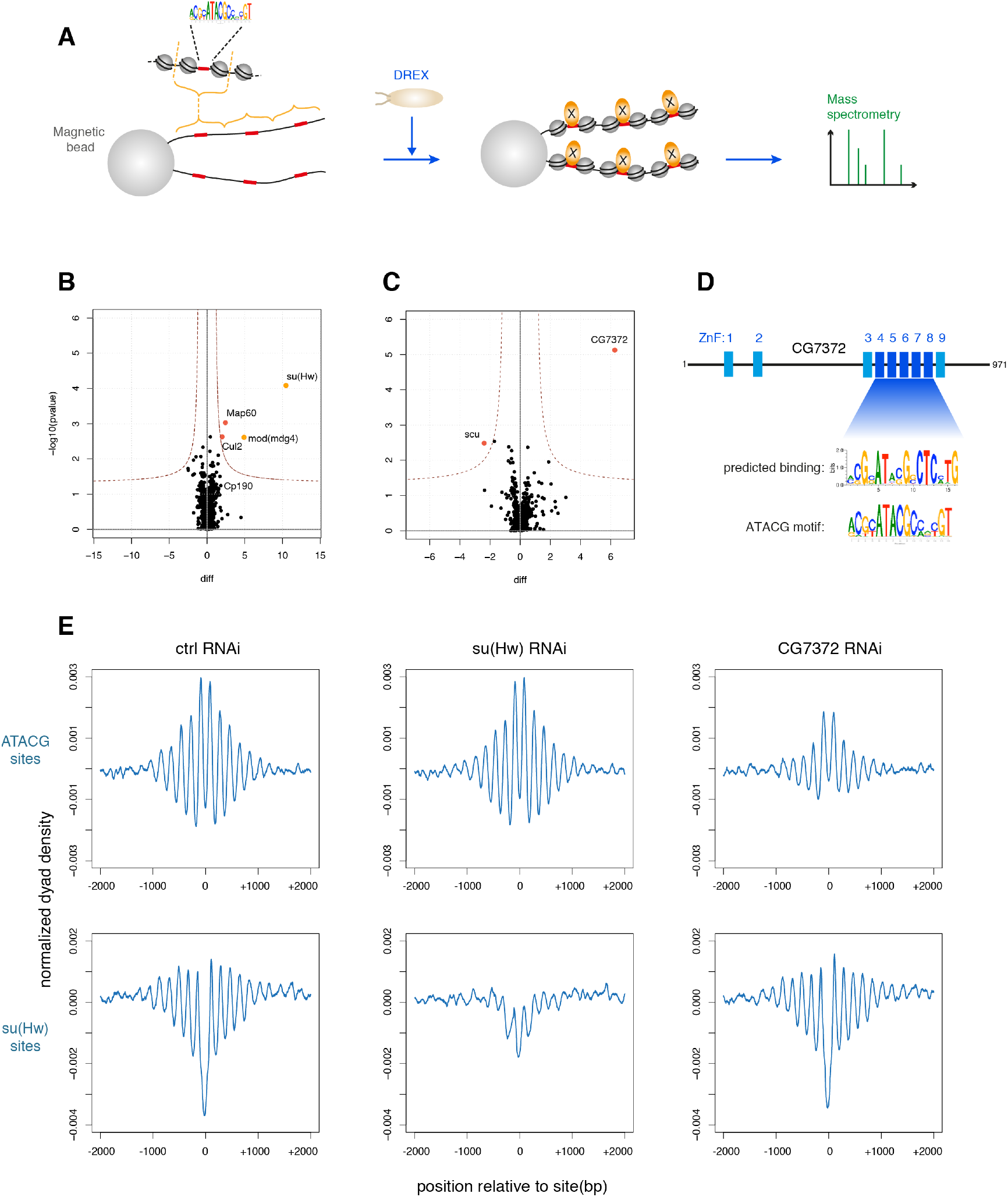
Identification of the ATACG-binding protein Phaser. (A) Adaption of the *in vitro* chromatin assembly assay to purify and identify DNA-binding proteins in their chromatin context. (B) Volcano plot of proteins enriched on chromatinized oligomers containing su(Hw) binding sites compared to the same oligomers with mutated su(Hw) sites. (C) Volcano plot of proteins enriched on chromatinized oligomers containing ATACG sites compared to the same oligomers with mutated ATACG sites. (D) CG7372 is an uncharacterized protein with multiple zinc finger domains. The computationally predicted binding motif for zinc fingers 3-9 closely resembles the ATACG motif. (E) Averaged nucleosome dyad densities aligned at ATACG (top) and su(Hw) motifs (bottom) in BG3-c2 cells depleted for su(Hw) or CG7372 by RNA interference.

As a positive control for the approach we attempted to purify known proteins associated with the su(Hw) motif. We cloned and oligomerized a 458 bp region around a genomic su(Hw) site that earlier had proven to efficiently assemble a PNA *in vitro.* We also generated an analogous DNA affinity construct with a mutated su(Hw) motif. Both arrays were coupled to paramagnetic beads, incubated in preblastoderm extract to assemble chromatin, and chromatin proteins were purified and analyzed by mass spectrometry (**Fig. 4B**). su(Hw) was indeed the most enriched protein on chromatin containing the su(Hw) binding motif, but we identified also mod(mdg4), a core component of the *gypsy* insulator (Gerasimova et al., 1995), and Map60, a known interactor of the insulator protein CP190 (Maksimenko et al., 2015) (**Fig. 4B**). This result validated the affinity purification strategy.

We then performed a similar chromatin assembly and purification assay with an ATACG-containing array. Remarkably, only a single protein was highly and significantly enriched compared to the mutant control (**Fig. 4C**): an as yet uncharacterized protein encoded by the gene *CG7372.* Interestingly, the protein is predicted to contain several zinc finger domains, which are well-described DNA binding modules (**Fig. 4D**). We then used a zinc finger target prediction tool that, based on the provided amino acid sequence, predicts the putative binding motif on the target DNA (Persikov and Singh, 2014). Strikingly, when zinc finger domains 4-8 of CG7372 are queried in this way, the predicted target motif very closely resembles the ATACG motif we identified at non-TSS PNAs (**Fig. 4D**). This strongly suggested that CG7372 was indeed the protein binding to ATACG sites *in vivo* and thereby responsible for nucleosome phasing around the motif. Therefore, we tested the effect of CG7372 knockdown by RNA interference on PNA formation in *Drosophila* BG3-c2 cells (**Fig. 4D**). When CG7372 was knocked down, regular phasing at ATACG sites was markedly reduced, while the PNAs around su(Hw) binding sites were not affected. Depletion of su(Hw), on the other hand, reduced phasing in the vicinity of su(Hw) sites but not at ATACG sites. These results suggest that CG7372 binds ATACG motifs *in vivo*, and, by positioning the −1 and +1 nucleosomes, sets up phased regular nucleosome arrays. CG7372 is, as of yet, an undescribed protein. We decided to name it Phaser (Phs), for its ability to strongly phase nucleosomes and apparently being one of the major determinants of nucleosome phasing throughout the genome.

### The CHRAC/ACF chromatin remodeler contributes to the formation of phased regular nucleosome arrays at ATACG sites

According to prevalent models, nucleosome phasing is achieved by the combined activity of barrier elements such as proteins that bind DNA tightly, and ATP-dependent nucleosome remodeling factors that slide nucleosomes against the barrier (Hartley and Madhani, 2009; Lieleg et al., 2015; Struhl and Segal, 2013; Wiechens et al., 2016; Zhang et al., 2011). We have identified Phs and su(Hw) as important barrier proteins for the formation of phased arrays. Next, we wanted to find which nucleosome remodelers may be involved in setting up phased arrays around Phs and su(Hw) binding sites. Dominant remodelers in preblastoderm embryo extracts are the Imitation SWItch (ISWI) family of remodelers CHRAC, ACF and NURF, which were first identified in this system (Ito et al., 1997; Tsukiyama and Wu, 1995; Varga-Weisz et al., 1997). Among those, the remodelling activity of NURF destroys regularity (Tsukiyama and Wu, 1995), but CHRAC and ACF complexes are prime candidates for contributing to PNA formation, since they promote the formation of regular nucleosome arrays both *in vitro* and *in vivo* (Fyodorov et al., 2004; Ito et al., 1997; Scacchetti et al., 2018; Varga-Weisz et al., 1997). To test the influence of CHRAC/ACF on array formation at ATACG sites, we obtained genome-wide nucleosome dyad maps for wildtype and *Acf*^7^ mutant embryos, which do not express Acf (Börner et al., 2016), the signature subunit of CHRAC/ACF (Ito et al., 1997; Varga-Weisz et al., 1997). Aligning the nucleosome dyads at ATACG sites revealed that regular phasing around the motif is strongly reduced in the *Acf* mutant (**Fig. 5A**). We found a similar pattern around su(Hw) motifs, although here the nucleosomes immediately flanking the sites are still well positioned in the mutant. The nucleosomes at the 5’ end of active of genes, on the other hand, do not show any discernible difference between wildtype and mutant embryos, suggesting that neither CHRAC nor ACF contribute to nucleosome phasing at TSS.

**Fig. 5.**
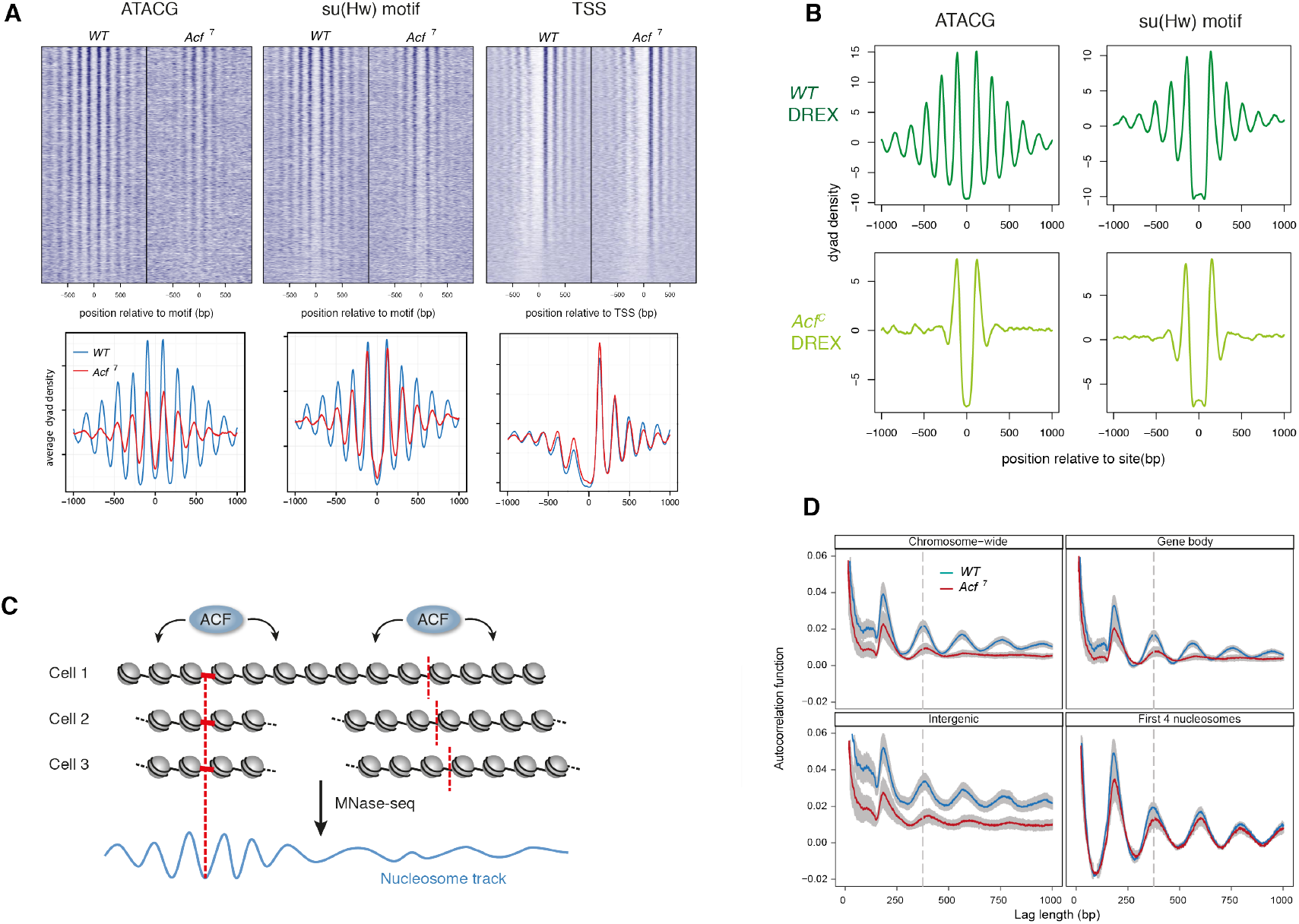
The CHRAC/ACF chromatin remodeler contributes to the formation of phased regular nucleosome arrays at ATACG sites. (A) Heat maps (top) and averaged profiles (bottom) of nucleosomal dyad densities aligned at ATACG sites, su(Hw) consensus motifs, and TSS in *wt* and *Acf^7^* mutant embryos. (B) Averaged nucleosomal dyad densities of chromatin assembled on genomic DNA with *wt* (top) or *Acf* DREX (bottom) aligned on ATACG and su(Hw) sites. (C) Illustration how regular nucleosome spacing promoted by a globally active CHRAC/ACF remodeler can only be appreciated in a mononucleosome map at sites with well-positioned nucleosomes (D) Global changes in chromatin regularity on chromosomes 2 and 3 are estimated using autocorrelation analysis (top left). The mean and SEM of replicate samples are displayed. The same analysis was performed for concatemerized gene bodies (top right), intergenic regions (bottom left), and 5’ end of genes (bottom right).

To explore whether the action of Acf-containing nucleosome sliding factors are observable in reconstituted chromatin we generated DREX from *Acf*^C^ embryos (where the complete Acf gene is deleted by CRISPR/Cas9 (Scacchetti et al., 2018)) and used it to assemble chromatin on genomic DNA. Mapping nucleosomes and alignment of dyad densities at the ATACG and su(Hw) sites revealed that the positioning of the two nucleosomes directly adjacent to the motifs was very similar in the mutant and the wild type extract (**Fig. 5B**). The regular arrays of flanking nucleosomes, however, were completely lost in chromatin assembled with *acf* mutant extract. Visual inspection of nucleosome tracks at single ATACG sites suggested that nucleosomes were still present in these flanking regions (**Supp Fig. 3**), but the spacing relative to the central nucleosomes is not constant anymore and the nucleosome dyad densities at these positions cancel each other out in the composite plot. Importantly, the loss of regularity around ATACG and su(Hw) sites in *Acf*^C^ mutant extract could be rescued by the addition of recombinant ACF remodeler (**Supp. Fig. 4**). In summary, our data suggest that Acf-containing complexes are not involved in positioning the nucleosomes directly adjacent to the barrier, but their nucleosome spacing activity aligns the outer nucleosomes relative to the central ones. Of note, the situation *in vivo* is less severe, possibly due to the activity of other remodelers with redundant functions during the later stages of embryogenesis (**Fig. 5A**).

### CHRAC/ACF has a global role in the formation of regular chromatin

Above, we have established that Acf was not required for positioning the central nucleosomes next to the ATACG and su(Hw) motifs, but rather for setting up regularly spaced arrays around them. In principle, there are two possibilities how this could be achieved. Either, CHRAC/ACF is somehow locally recruited to these sites to promote the formation of regular chromatin, akin to the specialized action of other remodelers at promoters. Alternatively, CHRAC/ACF might have a more global role in setting up regular spacing as has been suggested before (Fyodorov et al., 2004; Scacchetti et al., 2018). In the latter scenario, the *Acf* deficiency might affect the regularity of nucleosome arrays throughout the genome, but in the context of our study would only be appreciated at select sites with well-positioned nucleosomes, as regions with poorly positioned nucleosomes cannot be assayed for chromatin regularity in an MNase-seq experiment (**Fig. 5C**). As we had shown previously that CHRAC/ACF is required for setting up the repressive ground state of chromatin throughout the genome (Scacchetti et al., 2018), we favored the model of a global role of the remodeler.

To test whether loss of Acf affects chromatin regularity on a global level, autocorrelation function analysis was applied to mononucleosome maps from wild type and *Acf*^7^ mutant embryos. Autocorrelation analysis allows estimating the overall regularity of a continuous signal such as a nucleosome dyad density map (Braunschweig et al., 2009). The longer it takes for the autocorrelation function to dampen to a flat line, the more regular the underlying chromatin is. In a genome-wide analysis comprising chromosomes 2 and 3 (**Fig. 5D**) a significant dampening of the autocorrelation amplitudes was observed in the Acf^7^ mutant, which revealed that chromatin is less regular along the entire length of the mappable chromosomes in the absence of the remodeler. This demonstrates that Acf function is not confined to a few selected sites, but rather acts on a more global scale.

We also calculated the autocorrelation coefficients selectively for gene bodies, intergenic regions, and for the first 4 nucleosomes downstream of TSS (**Fig. 5D**). In these cases, dyad density profiles along the respective genomic regions were digitally concatemerized head to tail. Within gene bodies and intergenic regions, Acf depletion caused a decay of regularity as it did for the entire genome. On the other hand, overall regularity was not affected downstream of TSS in the *Acf*^7^ mutant. This suggests that, while Acf contributes to the bulk of nucleosome spacing in the genome, it does not affect the phasing at TSS, where other remodelers are known to be involved (Gkikopoulos et al., 2011b; Krietenstein et al., 2016; Yen et al., 2012). This also fits well with our observation that nucleosome phasing is not affected downstream of TSS in the *Acf*^7^ mutant when measured by metagene analysis (**Fig. 5A**). These results demonstrate that the CHRAC/ACF remodelers – through their global nucleosome spacing activity - are important to set up regularly phased arrays around Phs and su(Hw) sites.

## Discussion

### Spectral density estimation as a powerful tool for the genome-wide discovery of phased regular arrays

Traditionally, nucleosome phasing has been studied at well-established sites with strongly positioned nucleosomes. The phased nucleosome array downstream of the nucleosome-free regions at many active promoters is an important element of transcription control, and accordingly installed by the interplay between several chromatin remodelers and transcription factors (Hartley and Madhani, 2009; Krietenstein et al., 2016; Lieleg et al., 2015; Struhl and Segal, 2013; Zhang et al., 2011). It has been unclear to which extent other sites with similarly phased arrays exist, since the established approach of finding such nucleosome arrangements requires prior knowledge about candidate sites. Spectral density estimation (SDE) allows analyzing chromatin regularity in an unbiased manner throughout the whole mappable genome. Furthermore, by comparing the wild type SDE pattern with patterns derived from cells lacking chromatin components of interest, it should be possible to obtain sites that differ in regularity between the two conditions, and, thereby, to map potential target sites. Differential SDE maps between cell types or developmental stages could also indicate the location of important regulatory sites.

We applied SDE to the *Drosophila* genome to comprehensively identify locally phased regular nucleosome arrays (PNAs). The most abundant phasing element in the fly genome outside of promoters consists of a novel ATACG DNA motif. The protein Phs that binds to this element acts as a nucleosome phasing barrier at thousands of sites throughout the genome. Remarkably, we found this new factor not by the appearance of its footprint in the nucleosomal map or by an increase of DNA accessibility at target sites, but rather by its ability to locally position nucleosomes and to nucleate the formation of phased regular arrays. In fact, ATACG sites do not have increased accessibility, indicating that binding of Phs to its target motif does only obstruct a minimal stretch of DNA from being incorporated into nucleosomes. In the *Drosophila* genome, Phs appears to be the dominant barrier protein. The application of our strategy to other genomes may lead to the discovery of new chromatin proteins with similar properties.

### Chromatin assembly by *Drosophila* embryo extract for *in vitro* genomics and the purification of genomic features

Our second major technical innovation was the use of *Drosophila* preblastoderm embryo extract (DREX) to assemble and study chromatin *in vitro* on a genome-wide level. This physiological chromatin assembly system has been established some 25 years ago (Becker and Wu, 1992), but so far, it has been used mostly on biologically inert bacterial DNA. DREX-assembled chromatin is dynamic, containing abundant nucleosome sliding activities. Indeed, the paradigmatic ISWI-containing nucleosome sliding activities NURF, CHRAC and ACF have been first identified in this system (Ito et al., 1997; Tsukiyama and Wu, 1995; Varga-Weisz et al., 1997). Remarkably, the assembly of the entire *Drosophila* genome into chromatin using the DREX system not only recapitulates global features of physiological *Drosophila* chromatin, such as regular nucleosome spacing, but also the formation of PNAs. Focusing on the prominent PNAs around ATACG and su(Hw) sites, we purified, as a proof of principle, the associated proteins using chromatin affinity chromatography. In important extension of classical DNA affinity chromatography (Kadonaga and Tjian, 1986), we introduce chromatin affinity chromatography including nucleosomes flanking the presumed protein binding site.

Chromatin reconstituted in DREX is complex, containing several hundred chromatin components (Volker-Albert et al., 2016). Clearly, the genome-wide reconstitution of such physiological chromatin provides opportunities to study principles of chromatin organization beyond nucleosome phasing, such as transcription factor binding, chromatin accessibility, or the interaction of genomic regions. The assembled chromatin reflects the state of preblastoderm embryos before general transcription activation and the stratification of chromatin that characterizes differentiated cells. This preblastoderm chromatin may be seen and used as a ‘white page’ – a basal complex chromatin infrastructure that forms the foundation for more varied transcriptional landscapes (Blythe and Wieschaus, 2016). Nuclear extracts of older *Drosophila* embryos, on the other hand, have a high transcriptional activity (Kamakaka and Kadonaga, 1994; Robinson and Kadonaga, 1998) and may be added to study transcription of chromatin at a global scale. The approach could also be extended to other systems. In *Xenopus*, extracts from oocytes and eggs assemble chromatin on DNA and are transcriptionally active (Almouzni and Mechali, 1988; Laskey et al., 1977). *Xenopus* extracts may also be used to study the influence of the cell cycle on chromatin structure. For mammalian cells, there are well-established nuclear transcription extracts (Dignam et al., 1983). Chromatin may be reconstituted in advance by salt gradient dialysis or by replication-dependent chromatin assembly in nuclear extract (Krude and Knippers, 1993). Lastly, such *in vitro* genomic assays may also be performed in a biochemically fully defined environment, where, instead of extracts, purified components are used, as was pioneered in yeast (Krietenstein et al., 2016). Clearly, genome-wide chromatin biochemistry bears great potential for the mechanistic analysis of chromatin structure and function.

### The nature of phased regular sites throughout the genome

A good proportion of PNAs in the fly genome coincide with the 5’ ends of active genes, as expected. Interestingly, the remaining sites are much harder to connect to obvious genomic features and biological functions. We did not detect any significant overlap with enhancers, topological domain boundaries, or mapped histone modifications. However, a set of DNA motifs was found to be associated with regular sites. Several motifs correspond to sequence repeats of low complexity, which, as the salt-gradient nucleosome reconstitutions suggest, intrinsically promote or disfavor nucleosome formation and thus may lead to statistical positioning of nucleosomes in their immediate vicinity (Kornberg and Stryer, 1988). The generation of a locally confined nucleosome array is then due to the global nucleosome ‘spacing’ activity of chromatin remodelers such as CHRAC/ACF. It is possible that, at least for some of these sites, the occurrence of positioning sequences is by chance, and not connected to any biological activity.

Two prominent motifs localize in the centers of extended PNAs and function as binding sites for proteins. The su(Hw) protein that binds to one class of sites is a well-established constituent of the *gypsy* insulator complex, which is involved in insulating promoters from enhancer activity (Gause et al., 2001; Ghosh et al., 2001) and in the formation of topological-associated domains (Ramirez et al., 2018; Van Bortle et al., 2014). For Phs, which we identified due to its interaction with a new ATACG motif, a similar function is not yet known. It is somewhat surprising that we did not find a greater diversity of protein binding sites among the non-TSS PNAs. However, some of the proteins that are known to tightly bind to DNA and induce strong nucleosome positioning, such as the insulator protein CTCF, predominantly bind at promoters (Holohan et al., 2007) and therefore would not be picked up in this analysis.

### What is the biological function of ATACG sites and Phs?

Remarkably, PNAs containing the ATACG motif are not associated with prominent chromosomal features, such as promoters, enhancers, domain boundaries or certain histone modifications patterns. Conceivably, their major function could be within the unmappable part of the genome, such as constitutive heterochromatin and silenced repetitive elements, while the euchromatic sites may be just chance occurrences of the motif, unrelated to a function. However, the relative enrichment of the motif throughout the mappable genome argues for a positive selection of the motif. A very striking feature of the newly described ATACG motif is its strong ability to locally phase nucleosomes. Normally, only a fraction of the consensus motif sites of a particular DNA-binding protein is occupied in a cell. The observed strong phasing suggests that most of the sites are indeed tightly bound by Phs. Whether Phs contacts the two neighboring nucleosomes for more precise positioning remains to be tested. We also do not know whether regular nucleosome phasing around ATACG sites is required for Phs function or rather a side effect of strong protein binding. An interesting insight in the potential function of Phs comes from a study where hundreds of chromatin factors were systematically recruited to enhancers in *Drosophila* cells to assess their influence on the expression of a reporter gene (Stampfel et al., 2015). In this study, recruitment of Phs led to a significant reduction of transcription in many enhancer contexts, suggesting that the presence of Phs interferes with enhancer activation.

Interestingly, ATACG and su(Hw) sites share common features. Both bind multi-zinc-finger proteins, efficiently position adjacent nucleosomes, and are major determinants of nonpromoter PNAs. Even their core motifs are very similar (GCATACG vs GCATACT). However, su(Hw) binds to many target sites in a complex with the insulator proteins mod(mdg4) and CP190 (Schwartz et al., 2012). Accordingly, in our *in vitro* approach, we could copurify mod(mdg4) with su(Hw), but not with Phs. There is also no strong overlap between ATACG sites and published mod(mdg4) and CP190 binding profiles (Schwartz et al., 2012). This indicates that Phs is not part of a new insulator complex. It also fits with our observation that, unlike at su(Hw) sites, we do not observe an obvious expansion of the nucleosome linker bearing ATACG sites, arguing against the binding of a large multiprotein complex. Phs binding sites do not overlap with known functional elements, but, like su(Hw) sites (Sexton et al., 2012), are enriched in inactive chromatin domains. su(Hw) and other architectural proteins have been shown to guide chromosome architecture within chromatin domains (Rowley et al., 2017). Maybe Phs and su(Hw) contribute to chromosome architecture specifically within silent domains by similar mechanisms.

As the ATACG motif seems to promote phasing in mammals as well, it would be interesting to identify the mammalian protein binding to it. We could not detect any obvious mammalian homologue of Phs - however, the determination of orthologues of multi-zinc-finger proteins is notoriously difficult.

### Global effects of chromatin remodeling may only be detected at localized sites

We identified CHRAC/ACF as a remodeler required for aligning regularly spaced nucleosomes adjacent to ATACG and su(Hw) sites. However, the activity of CHRAC/ACF is not confined to these regions, but it has a more global role promoting chromatin regularity in a genome-wide manner. Chromatin regularity is defined by an even spacing between the nucleosomes, but does not require that the nucleosomes are well-positioned relative to the underlying DNA-sequence. This means that a particular chromatin region may be very regular in all the cells of the population, but because the nucleosomes are differently positioned in different cells, the regularity cannot be appreciated in a population-based mononucleosomal map (compare **Fig 5C**). CHRAC/ACF has been shown to sense the linker length between nucleosomes in an array *in vitro* and shift and slide nucleosomes until there is an even spacing between all of them (Hwang et al., 2014; Yang et al., 2006). It does, however, not account for the positioning of the nucleosomes relative to the underlying DNA sequence. Phased nucleosome arrays originate from the interplay between barrier elements and the general nucleosome spacing activity of nucleosome sliding factors. Barrier elements may position adjacent nucleosomes due to their intrinsic DNA sequence properties or due to tightly bound protein. The positioning of the next nucleosome neighbors may be tightly regulated, such as at promoters, or statistically following the tendency of the system for maximal charge neutralization of the DNA. Further phasing of an array of nucleosomes then only requires a general nucleosome spacing activity provided by CHRAC/ACF (Fig. 6). The global role of the remodeler, which we detected by autocorrelation analysis, is best appreciated at localized sites with strongly positioned nucleosomes.

**Fig. 6.**
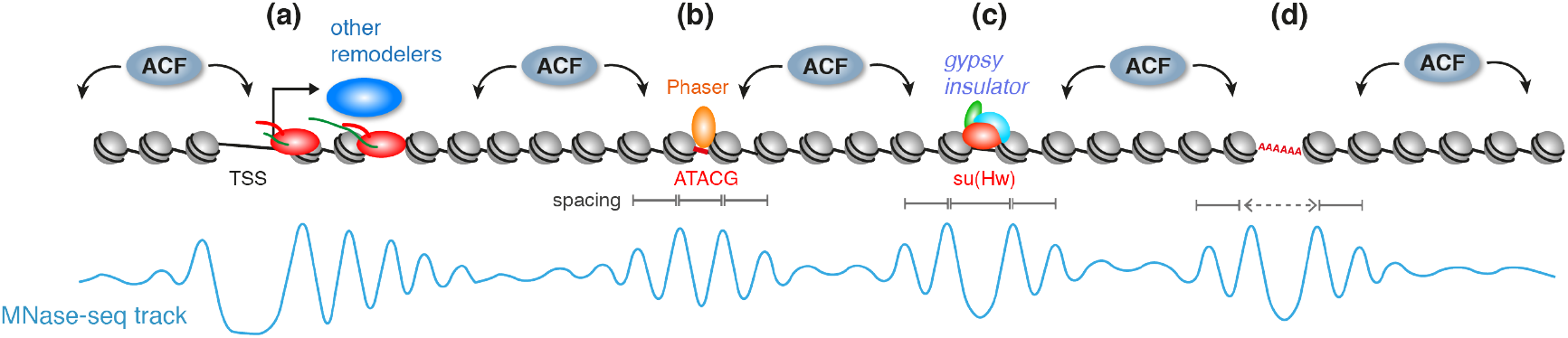
Model. Phased nucleosome arrays originate from the interplay between barrier elements and the general nucleosome spacing activity of nucleosome sliding factors. At promoters, nucleosome-free regions and the binding of general transcription elements, together with dedicated chromatin remodelers and the transcriptional machinery, establish regular arrays downstream of the TSS (a). Outside of promoters, chromatin regularity is established by the global activity of the CHRAC/ACF remodeler. However, this can only be appreciated at select sites with well-positioned nucleosomes. Major punctuation elements that locally position adjacent nucleosomes are Phaser binding sites (b), *gypsy* insulator sites (c), and DNA sequences with intrinsic nucleosome positioning properties (d). Importantly, the formation of locally confined, phased regular arrays does not require any specific recruitment of chromatin remodeling enzymes to these sites.

Importantly, we could not detect any effect of CHRAC/ACF on PNA formation at the 5’ end of genes, presumably because these critical arrays are installed by several other nucleosome remodeling factors dedicated to transcription control (Gkikopoulos et al., 2011b; Krietenstein et al., 2016; Yen et al., 2012).

If CHRAC/ACF has indeed this global role in providing chromatin regularity, then what are its biological functions? *Acf* mutant flies are viable, but show an increased embryonic lethality (Fyodorov et al., 2004). We recently found that the regularity of nucleosome spacing installed by CHRAC/ACF contributes to the repressive ground state of chromatin (Scacchetti et al., 2018). This function fits well to our observation that the chromatin in *Acf* mutant cells loses much of its regularity and therefore may become ‘leaky’ to untimely transcription at a global scale.

## Methods

### *Drosophila* strains

Two wild type genotypes (*w1118, yw*) were used to establish reference maps of nucleosome positions. The *Acf*^3^ mutant contains a 3098 bp deletion in the *Acf* gene from the first intron to the third exon (Börner et al., 2016). The *Acf^C^* mutant has most of the coding region deleted and is described in (Scacchetti et al., 2018).

### Nucleosome mapping in embryos

For mapping nucleosomes in *Drosophila* embryos were collected 2-8 hr after egg laying. Briefly, eggs laid during a 6 hr period were collected and aged for a further two hours at 25°C. Subsequently, one gram embryos were dechorionated in 120 ml 1:5 diluted sodium hypochloride for 3 min, thoroughly washed and fixed in 10 ml fixing solution (0.1 M NaCl, 0.05 M HEPES pH 8.0, 1 mM EDTA, 0.5 mM EGTA, 1.8% Formaldehyde added to 30 ml n-Heptane) for 15 min at 16-18°C on a rotating wheel. Fixation was quenched by adding 125 mM f.c. glycine. The embryos were subsequently washed with PBS (including 0.01% Triton-X100) for 10 min and stored at −80°C.

For nuclei isolation, embryos were slowly thawed and dounced using a glass homogenizer (Schubert, Cat.no. 9164693) with 20 strokes each of the A and B pestles in ice-cold NX-I buffer (15 mM HEPES pH 7.6, 10 mM KCl, 2 mM MgCl2, 0.5 mM EGTA, 0.1 mM EDTA, 350 mM sucrose, 1 mM DTT, 0.2 mM PMSF, Protease inhibitors Leupeptin, Pepstatin and Aprotinin (10 μg/ml)). The lysate was filtered through Miracloth and nuclei were pelleted at 3,500 rpm, 10 min at 4°C.

For MNase digestion, the nuclei were resuspended in 1 ml MNase digestion buffer (10 mM Tris-HCl pH 7.5, 15 mM NaCl, 60 mM KCl, 2 mM CaCl_2_, 0.15 mM spermine, 0.5 mM spermidine). About 80 mg of nuclei were digested with 0.15 U of MNase (Sigma, Cat.no N5386) for 15 min at 37°C while shaking at 500 rpm to yield predominantly mononucleosomes. The reaction was stopped by adding 20 μl of 0.5 M EDTA, pH 8.0 and the tubes were quickly transferred to ice for 1 min. Nuclei were spun at 12,500 rpm, 10 min at 4°C. The supernatant containing most of the DNA was collected and the buffer composition was adjusted to RIPA (1% Triton X-100, 0.1% Na deoxycholate, 140 mM NaCl, 10 mM Tris pH 8.0, 1 mM EDTA, 0.1% SDS, 1mM PMSF). The chromatin was sheared in a Covaris sonicator in a 12×12 fiber tube at 50 W, 20% output for 8 min. After a centrifugation at 17,000 × g for 20 min, the supernatant was used for subsequent H3 ChIP.

Chromatin immunoprecipitation (ChIP) was performed by first pre-clearing the chromatin sample using a protein A+G (1:1) bead mix for 1 hr at 4°C. Then, 150 μl of chromatin was incubated overnight with 3.5 μl rabbit polyclonal anti-H3 antibody (Abcam, Cat.no ab1791) in a final volume of 500 μl. Nucleosomes were immunoprecipitated by adding a (1:1) mix of protein A+G beads for 3 hr at 4°C. Subsequently, the beads were washed 5 times in 1 ml RIPA buffer. Residual RNA was digested by RNase-A (10 μg/100 μl, Sigma, Cat. No. R4875) at 37°C for 20 min. Subsequent protein digestion (using 25 μg/100 μl of Proteinase K) and reversal of crosslinking were performed simultaneously at 68°C for 2 hr. DNA was purified using GenElute™ PCR Clean-Up Kit (Sigma).

Recovered DNA was quantified using the Qubit® dsDNA HS Assay Kit (Life Technologies) and sequencing libraries were prepared using the MicroPlex Library Preparation kit (Diagenode) starting with 2 ng DNA whenever possible. PCR amplification was monitored by quantifying amplified libraries (maximum 19 cycles). The libraries were sequenced on a HighSeq 1500 (Illumina) instrument.

### RNAi and nucleosome mapping in cultured cells

BG3-c2 cells were cultured at 26°C in Schneider’s *Drosophila* Medium (Gibco) including 10% fetal calf serum (FCS), Penicillin-Streptomycin and 10 μg/ml human insulin. For knockdown of su(Hw) and CG7372 two different dsRNAs per gene were simultaneously used. For each knockdown condition, 1.5 × 10^7^ cells were resuspended in 4 ml medium (containing antibiotics and insulin but no FCS), seeded in T-75 flask, and incubated with 100 μg (GST, su(Hw), CG7372) of *in vitro* transcribed dsRNA (MEGAscript T7 Transcription Kit, Ambion; primers in Supp. Table 3). After gentle shaking for 45 min at room temperature, 16 ml of medium (+FCS/+antibiotics/+insulin) was added to the cells. After 5 days cells were treated again with dsRNA in the same manner, but this time two T-75 flasks per condition were seeded. After 5 more days, the cells were harvested. 9 × 10^7^ cells per condition were washed in 10 ml cold PBS and resuspended in 1 ml TM2 buffer (10 mM Tris-HCl pH 7.5, 2 mM MgCl_2_, 1 mM PMSF). The cells were incubated 1 min on ice and NP-40 added to 1.5 %, gently vortexed, and again incubated on ice for 10 min for lysis and release of the nuclei. After a 10 min centrifugation at 93 × g and 4°C the nuclei were resuspended in 600 μl of TM2 buffer with 2 mM CaCl2 and distributed into three 200 μl aliquots. Those were treated with 2.3/5.9/11.7 × 10^−3^ units of MNase for 20 min at 37°C. The reaction was then quenched by adding 5 μl 0.5 M EDTA and 100 μg RNase A, followed by a 20 min incubation at 37°C. Proteins were digested away at 50°C for 2 h after the addition of 12 μl 10% SDS and 200 μg Proteinase K. The DNA was then purified by Phenol/Chloroform extraction, precipitated in 2.5 V EtOH, 1/10 V NaOAC 3M and 20 μg glycogen and run on a 1.2% agarose gel in 0.5x TAE buffer. The samples with the best digestion degree (predominantly nucleosomes, achieved with 11.7 × 10^−3^ U of MNase) were chosen and the mononucleosomal bands cut out and purified (NucleoSpin Gel and PCR Clean-up, Macherey-Nagel). The purified DNA was then used for Next-Generation-Sequencing library preparation.

### Nucleosome position data analysis

Raw paired-end reads were mapped to the *D. melanogaster* dm3 genome assembly with Bowtie v1.1.1 (Langmead et al., 2009). Dyad coverage vectors were obtained by size-selecting fragments of length >=120 and <=200 bp and resizing their length to 50 bp fixed at the fragment center.

### Spectral density estimation

For identifying regions of regular nucleosome phasing, the nucleosome signal along each chromosome was scanned in a sliding window of 1024 bp with a step of size 100 bp. In each window a periodogram was obtained using the R function spec.pgram with parameters log=“n”, pad=1, spans=c(3,3). The spectral density corresponding to a 192 bp period was extracted and log-transformed. Phased nucleosome arrays were identified by averaging z-score transformed spectral densities of all wildtype samples. Regions passing an arbitrary threshold of 2.5 were defined as phased nucleosome arrays (PNA).

### Autocorrelation analysis

The autocorrelation function was calculated for the dyad coverage vectors obtained for the entire genome, gene bodies, first 4 nucleosomes or intergenic regions in the R environment. The vectors for the last 3 cases represent head-to-tail concatemerized regions of given annotation considering their orientation. The function was run for the lag length of 1000 bp.

### Motif analysis

We searched for enriched motifs in PNA regions using MEME (MEME suite version 4.10.0) (Bailey et al. 2009) using the zero or one occurrence per sequence (“zoops”). Genome-wide searches for motif hits were performed with FIMO using standard settings. The top 5 % of motif hits were chosen for further analysis. Instances of the motifs within non-TSS PNA regions were found with MAST using standard settings.

### Drosophila embryo extract (DREX) preparation

Drosophila embryo extract was prepared from preblastoderm embryos 0-90 min AEL according to (Becker and Wu, 1992) with the following modifications: After collection, the embryos were dechorionated in 200 ml embryo wash buffer (0.7% NaCl, 0.04% Triton X-100) and 60 ml 13% sodium hypochlorite (VWR) for 3 min at room temperature while stirring. Embryos were rinsed for 5 min with cold water and transferred into a glass cylinder with embryo wash buffer. After settling of the embryos, the buffer was decanted and the embryos were washed first in 0.7% NaCl and then in extract buffer (10 mM HEPES, pH7.6, 10 mM KCI, 1.5 mM MgCl_2_, 0.5mM EGTA, 10% glycerol, 10 mM 3-glycerophosphate; 1 mM DTT, protease inhibitors added freshly before use: 0.2 mM PMSF, 1 mM Aprotinin, 1 mM Leupeptin, 1 mM Pepstatin). After the last wash, embryo wash buffer was decanted and embryos were homogenized with one stroke at 3000 rpm and 10 strokes at 1500 rpm using a homgen^plus^ homogenizer (Schuett-Biotec). The homogenate was adjusted to a final MgCl2 concentration of 5 mM and centrifuged for 15 min at 27000 g at 4°C. The white lipid layer was discarded and the supernatant was centrifuged for 2 h at 245420 g at 4°C. The clear supernatant was aspirated with a needle, leaving the lipid layer and pelleted nuclei in the tube.

### DREX-mediated *in vitro* chromatin assembly

Genomic DNA was purified from *Drosophila* BG3-c2 cells with the Blood & Cell Culture Midi Kit (Qiagen). 1 μg of genomic or plasmid DNA were added to 60 μl *Drosophila* embryo extract supplied with 10 mM 3-glycerophosphate and an ATP-regenerating buffer (3 mM ATP, 30 mM creatine phosphate, 10 μg/ml creatine kinase, 3 mM MgCl_2_, 1 mM DTT) and filled up to a total volume of 120 μl with EX50 buffer (10 mM HEPES, pH 7.6, 50 mM KCI, 1.5 mM MgCl_2_, 0.5 mM EGTA, 10% glycerol, 10 mM 3-glycerophosphate; 1 mM DTT, 0.2 mM PMSF, 1 mM Aprotinin, 1 mM Leupeptin, 1 mM Pepstatin). Chromatin was assembled at 26°C for 6 h (for nucleosome mapping) or 1 h (for mass spectrometry). For the rescue of phasing in *acfΔ* mutant DREX by recombinant ACF, ACF was purified from insect cells as described in (Fyodorov and Kadonaga, 2003) and 2.4 pmol added to the chromatin assembly reaction.

For nucleosome mapping, 1.5 mM f.c. CaCl_2_ and 1.4 × 10^−3^ U MNase were added to digest the chromatin at 30°C for 15 min. After quenching the reaction with 10 mM f.c. EDTA, samples were treated with 20 RNase μg A for 30 min. Proteins were digested overnight after adding 16 μl 2% SDS and 100 μg Proteinase K and incubating at 37°C. The DNA was purified with the GenElute PCR clean-up kit (Sigma-Aldrich) and libraries for paired-end sequencing generated as described above.

### Chromatin assembly by salt-gradient dialysis

A 100 μl assembly reaction containing 10 μg genomic DNA, ~10 μg purified *Drosophila* histone octamers (gift from M. Walker and P. Korber, exact amount optimized by titration), 20 μg BSA, 0.1% Igepal CA-630, 10 mM Tris pH 7.6, 2 M NaCl, 1 mM EDTA were prepared and transferred to a dialysis cup (Slide-A-Lyzer; MWCO 3500, Thermo Fisher). The dialysis cup was then floated in a big beaker containing 300 ml high salt buffer (10 mM Tris-HCl pH 7.6, 2 M NaCl, 1 mM EDTA pH 8.0, 0.05% Igepal CA-630, 0.1% 2-mercaptoethanol). NaCl concentration in the beaker was decreased constantly at room temperature over night by pumping 3 l of low salt buffer (10 mM Tris pH 7.6, 50 mM NaCl, 1 mM EDTA, 0.05% Igepal CA-630, 0.01% 2-mercaptoethanol) into the beaker with a peristaltic pump (Minipulse evolution, Gilson, mode 8.4 rpm). After the entire low salt buffer had been transferred, the dialysis cup was dialyzed another 2 h at room temperature against fresh low salt buffer. The quality of nucleosome assembly was assessed by limited MNase digestion.

### Identification of motif-interacting proteins in DREX-assembled chromatin

DNA around motif sites was amplified by PCR (50 μl PCR per construct) from plasmids with cloned genomic regions (primers in Supp. Table 3), purified with the GenElute PCR clean-up kit and eluted in 50 μl. To phosphorylate the ends of the amplified fragments, they were incubated at 37°C for 2 h in phosphorylation conditions (1x T4 ligase buffer (NEB), 1.2% PEG6000, 10 mM DTT, 5 mM ATP, 25 U T4 Polynucleotide kinase (NEB)). Ligation for array formation was performed by adding 800 U T4 DNA ligase (NEB) and incubating at 4°C for 30 min and then at 16°C overnight. After verifying fragment oligomerization on an agarose gel, the ligated DNA was purified by phenol/chloroform extraction and ethanol precipitation and dissolved in 37 μl H_2_0. To biotinylate the ends, 5 μl 10x NEBuffer 2 (NEB), 5 μl 0.4 mM Biotin-14-dATP (Life Technologies) and 15 U Klenow (exo-) (NEB) were added, followed by an overnight incubation at 37°C. The biotinylated DNA was then purified with Quick Spin Columns for radiolabeled DNA purification (Sigma-Aldrich). To find proteins that would bind to su(Hw) or ATACG motifs, 4 replicates of arrays with unaltered motifs and 4 replicates of mutated arrays were used for chromatin assembly and subsequent mass spectrometry. For each replicate, 2 μg of biotinylated DNA were coupled to 15 μl of M280 Streptavidin Dynabeads (Life Technologies) at 4°C overnight. The beads had been blocked before at room temperature for 1 h in PBS with 0.05% BSA and 0.05% NP-40. Chromatin assembly with DREX was performed on 2 μg DNA on beads as described above (all volumes in the assembly were doubled to take into account the double amount of DNA). The assembled chromatin was washed once with EX50 buffer (w/o protease inhibitors) and then incubated in 20 μl NuPAGE LDS sample buffer (Invitrogen) at 70°C for 10 min to elute associated proteins. Eluted proteins were separated by gel electrophoresis using a 4-12% Bis-Tris gel in 1X MES buffer (Thermo) for 8 min at 180 V. Proteins were fixated, stained with coomassie brilliant blue and each gel lane was minced. Sample destaining, reduction and alkylation was followed by an overnight digestion with trypsin (Scheibe et al., 2013). Prior to mass spectrometry analysis peptides were extracted, concentrated and desalted (Rappsilber et al., 2007). The samples were injected via a 20 cm long capillary (75 μm inner diameter, New Objective), packed in-house with ReproSil-Pur 120 C18-AQ, 1.9 μm (Dr. Maisch) mounted in a column-oven (Sonation). Peptides were eluted with an 88 min gradient of 5-40 percent ACN (Easy-nLC 1000, Thermo). The QExactive Plus mass spectrometer (Thermo) was operated in a data-dependent acquisition strategy performing up to 10 MS/MS scans (17,500 resolution) per full scan (70k resolution, 30-1650 m/z) in the orbitrap analyzer. The acquired data was processed with the MaxQuant software suite (ver1.5.2.8;(Cox and Mann, 2008)) using standard settings with activated match between runs and LFQ quantitation (Cox et al. 2014) and the supplied Drosophila uniprot database (18,767 entries). The data was post-processed in the R framework (ver3.1.2) filtering for a minimum of 2 peptides with at least one of them unique and removing protein groups only identified by site, reverse hits and known contaminants. The median log2 transformed LFQ quantitation values were subtracted and plotted together with the −log10 p-value (welch t-test between bait and control).

## Acknowledgments

We thank Silke Krause for the purification of recombinant ACF, Stefan Krebs and the LAFUGA Genomics Facility for next generation sequencing and Raffaella Villa and Alessandro Scacchetti for critical reading of the manuscript. This work was supported by grants Be1140/8-1 and SFB-1064-A01 of the German Research Council (DFG) to PBB.

## Author Contributions

S.B., D.J., T. S., and P.B.B. conceived experiments; S.B., D.J., L.H., M.S. and A.Z. performed experiments; S.B., D.J., M.S., and T.S. analyzed data; F.B., T.S. and P.B.B. provided feedback and supervision, S.B. and P.B.B. wrote the manuscript, F.B. and P.B.B. secured funding.

